# Synchronization Temperature Determines the Location of RSV Fusion During Entry in Cultured Cells

**DOI:** 10.1101/2021.03.09.434545

**Authors:** Christian Cadena-Cruz, Marcio De-Avila-Arias, Heather M. Costello, Leidy Hurtado-Gomez, Walter Martínez-De-La-Rosa, Gigliola Macchia-Ceballos, Wendy Rosales-Rada, Gerardo Valencia-Villa, Pedro Villalba-Amarís, Meisam Naeimi Kararoudi, Mark E. Peeples, Homero San-Juan-Vergara

## Abstract

Respiratory syncytial virus (RSV) is the most frequent cause of bronchiolitis in children under five years of age. No vaccines against this virus are currently available. RSV infection of a cell is initiated by fusion between the virion membrane and a cellular membrane, but it is not clear if the fusion process takes place at the plasma membrane or within an endosome. Most such experiments have been initiated at the traditional synchronization temperature of 4°C, an abnormal temperature for animal cells and one at which cellular homeostasis may be negatively affected. We have compared two synchronization temperatures (4°C and 22°C) to determine the kinetics of RSV entry into human bronchial epithelial cells. Following inoculation, virus entry was halted at different times by the addition of neutralizing antibody or temperature reduction to 4°C. We engineered a virus that encodes an extra viral gene, beta-lactamase fused to the viral phosphoprotein (P), to enable rapid detection after infection initiation. We found that the synchronization temperature used during inoculation determines the site of fusion. Transition from 4°C to 37°C resulted in RSV entry via the endosomal pathway but also induced F-actin disruption and plasma membrane blebbing, whether the cells were inoculated with RSV or not. Transition from 22°C to 37°C resulted in RSV entry by fusion at the plasma membrane and without the F-actin and plasma membrane disruptions. These results suggest that RSV normally enters cells by fusion at the plasma membrane and that the induction of endocytosis by infection synchronization at 4°C may be an artefact caused by distortion of the plasma membrane-supporting cytoskeleton.

**Author Summary:** In order to understand the overall mechanism driving infection, it is important to determine how the virus enters cells. The pathway that RSV uses to infect cells is unclear. It is a common practice to attach the virions at 4°C to synchronize the viral infection. In this report, we found that warming up primary cultures of undifferentiated normal human bronchial epithelial cells to 37°C from 4°C triggered dramatic changes in their cell membrane and cytoskeleton totally unrelated to the presence of the virions. The assessment of viral content delivery to the cytoplasm using RSV engineered to express BlaM allowed us to find that the virions attached at 4°C or 22°C fused their envelope with endosome or plasma membrane, respectively. Consequently, the entry via endosome after attachment at 4°C is an experimental artefact and RSV infects by fusing its envelope with the plasma membrane. The implications go beyond RSV since the entry of several virus species have been explored by synchronizing the infection after attachment at 4°C.

## Introduction

Respiratory syncytial virus (RSV), a member of the *Pneumoviridae* family of non-segmented negative sense RNA viruses [1], is a major contributor to acute lower respiratory tract infection in young children (bronchiolitis and pneumonia) and elders (pneumonia) [2]. In addition, RSV infection is one of the major causes of acute exacerbations of chronic obstructive pulmonary disease in older adults and an important contributor to morbidity and mortality in this group [3,4]

The RSV envelope is derived from the infected cell membrane and incorporates three virus-encoded proteins [5,6]. The G glycoprotein mediates the attachment process [6] and the F glycoprotein protein causes fusion between the virion membrane and the target-cell membrane [7]. The SH protein may act as a viroporin [8] and provides resistance to the action of tumor necrosis factor-alpha (TNF-α) [9]. Several proteins in the target cell membrane interact with either the G or F proteins. Annexin-2 [10] and heparan sulfate-rich proteoglycans [11] have been identified as receptors for the G protein in immortalized cells, but CX3CR1 appears to be a receptor for the G protein in primary airway epithelial cultures [12–15]. The F protein may also have a receptor and evidence has been presented for both nucleolin and ICAM-I in this role [16–18].

Once an RSV virion contacts a cell, it likely glides or tumbles over the cell plasma membrane as a consequence of either the G protein binding to charged molecules embedded in the cholesterol-rich membrane domains [19,20], but the location within or on the cell where the fusion process takes place is not clear. Srinivasakumar et al. provided evidence that the virus fuses with the plasma membrane [21]. However, Kolokoltsov et al., using siRNA to reduce expression of clathrin pathway-associated molecules, dramatically decreased RSV infection suggesting entry via an endocytic pathway [22]. Krzyzaniak et al. and San-Juan-Vergara et al, found that dequenching of fluorescently labeled virions took place in the endosomes pointing to macropinocytosis as the RSV entry mechanism [20,23].

It is possible that the temperature of virus adsorption influences the RSV entry mechanism. To test this possibility, we inoculated primary undifferentiated human bronchial epithelial (NHBE) cells [24] with RSV engineered to express the beta-lactamase (BlaM) reporter protein fused to the viral phosphoprotein (P), allowing rapid detection of the delivery of viral contents into the cytoplasm. Following the approach of Melikyan et al. [25], we determined whether virions preferentially fused their envelope with the plasma membrane or the endosomal membrane. Synchronization at 4°C triggered changes in the plasma membrane that facilitated endocytosis of cell membrane-adsorbed virions. The same experiment performed with a synchronization temperature of 22°C, resulted in RSV fusing its envelope with the plasma membrane. Consequently, the cell response to the non-physiological temperature of 4°C during viral attachment alters the mechanism by which the virus enters these cells.

## Results

### Incubation at 4°C disrupts actin filaments and induced cell blebbing

Attaching virus at 4°C for 1 h is the most common method for synchronizing virus entry and infection. We used F-actin status as a proxy to determine how incubation at different temperatures impacts cell physiology. Alexa-488-labeled phalloidin was added to visualize F-actin in NHBE cell cultures, which were incubated at 4°C, 22°C, and 37°C for 1 h. F-actin in cells incubated at 37°C was used as a reference for normal F-actin morphology (**Fig 1A**). In cells subjected to incubation at 4°C, there was a striking reduction in the thickness of the cortical actin region, stress fibers disappeared, F-actin accumulated in the perinuclear region, and small clumps of F-actin were scattered throughout the cytoplasm (**Fig 1B**). After warming from 4°C to 37°C for 10 min, actin clusters disappeared, stress fibers start reappearing and actin-based rings appeared at the plasma membrane in most of the cells. (**Fig 1C**). Between 40-60% of all cells (20-38 cell per field) assessed in 10 different fields showed actin-based rings. Such actin-based rings are associated with the blebbing phenomenon [26,27]. On the other hand, cells incubated at 22°C did not show significant changes in the F-actin morphology compared with that observed at 37°C (**Fig 1D**), and actin rings were not observed at the cell membrane after warming from 22°C to 37°C (**Fig 1E**). NHBE cells survive these conditions, but attaching the virus at 22°C, appears to be less disruptive to cell morphology and perhaps physiology.

**Fig 1.**
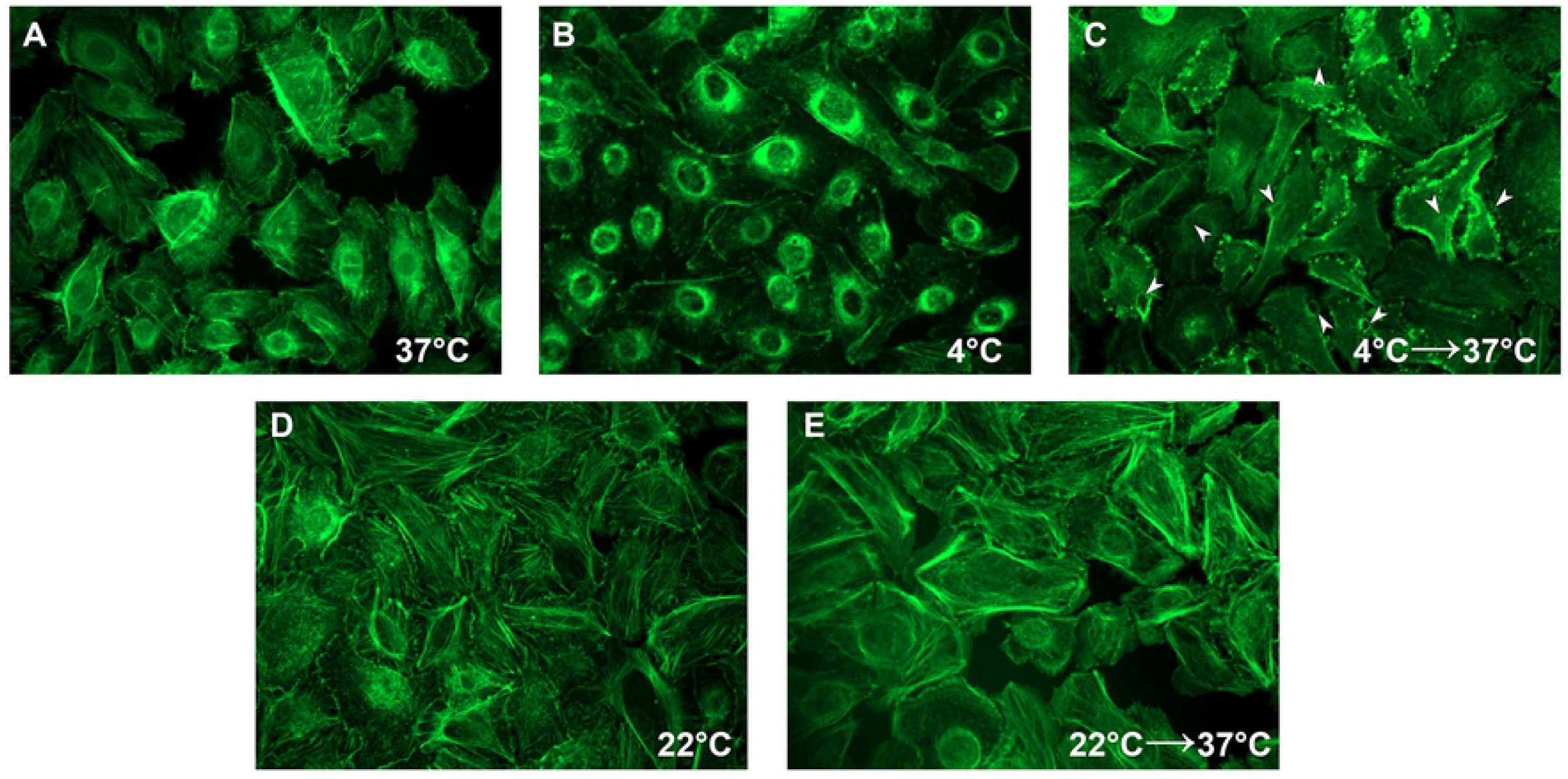
Cell membrane F-actin rearranges at 4°C and membrane blebbing occurs after a return to 37°C in a virus-independent manner. NHBE cells grown on poly-lysine-treated coverslips until they reached 70% confluency were incubated at the indicated temperature for 1 h, and followed by 10 min at 37°C where indicated (C, E). Cultures were fixed with 4% PFA, permeabilized and treated with Alexa 488-labelled Phalloidin to identify the position of F-actin.

### RSV uses different pathways to enter NHBE cells depending on their temperature during attachment

Miyauchi et al [25] adapted the approach developed by Cavrois [28] taking advantage of Beta-lactamase (BlaM) as a reporter to determine whether a virus enters via fusion at the plasma membrane or endocytosis. We determined the infection kinetics by using antibodies to neutralize virus entry at the plasma membrane or cold temperature (4°C) to universally hinder virus fusion with the plasma- and endosomal-membranes. Going forward, we will use “Antibody Block (AB)” and “Temperature Block (TB)”, respectively, to indicate the method used. Cold temperature reduces the lipid fluidity of the membranes. If the entry kinetics of both approaches coincide, the virus has fused with the plasma membrane. If the entry kinetics in TB are delayed (displaced to the right), the virus entered through endocytosis since it would take time for the virus to reach the endosome or compartment where the fusion takes place [25].

To express BlaM from rgRSV we inserted an additional gene encoding a fused P-BlaM protein immediately downstream of the P gene (**Fig 2A**). After virus recovery and growth, rgRSV-P-BlaM virions were pelleted by centrifugation, lysed, and shown to contain P-BlaM by immunoblotting with an anti-BlaM antibody (**Fig 2B**). Flow cytometry was used to evaluate the functional activity of BlaM through the cleavage of CCF2 in undifferentiated NHBE cells that were previously infected with rgRSV-P-BlaM (**Fig 2C**). The CCF2 accumulation in the cells and its concurrent cleavage took place at 17°C for 3 hr. The cleavage of a fraction or the totality of CCF2 by BlaM in the cytoplasm was revealed by shifting the emission from 520 nm to 440 nm after excitation at 405 nm due to the release of hydroxycoumarin (**Fig 2C**). The infectious titer of rgRSV-P-BlaM measured using BlaM activity after 3 hr matched the titer determined after 16 h using GFP fluorescence (**Fig 2C**) confirming the association of BlaM expression from rgRSV.

**Fig 2.**
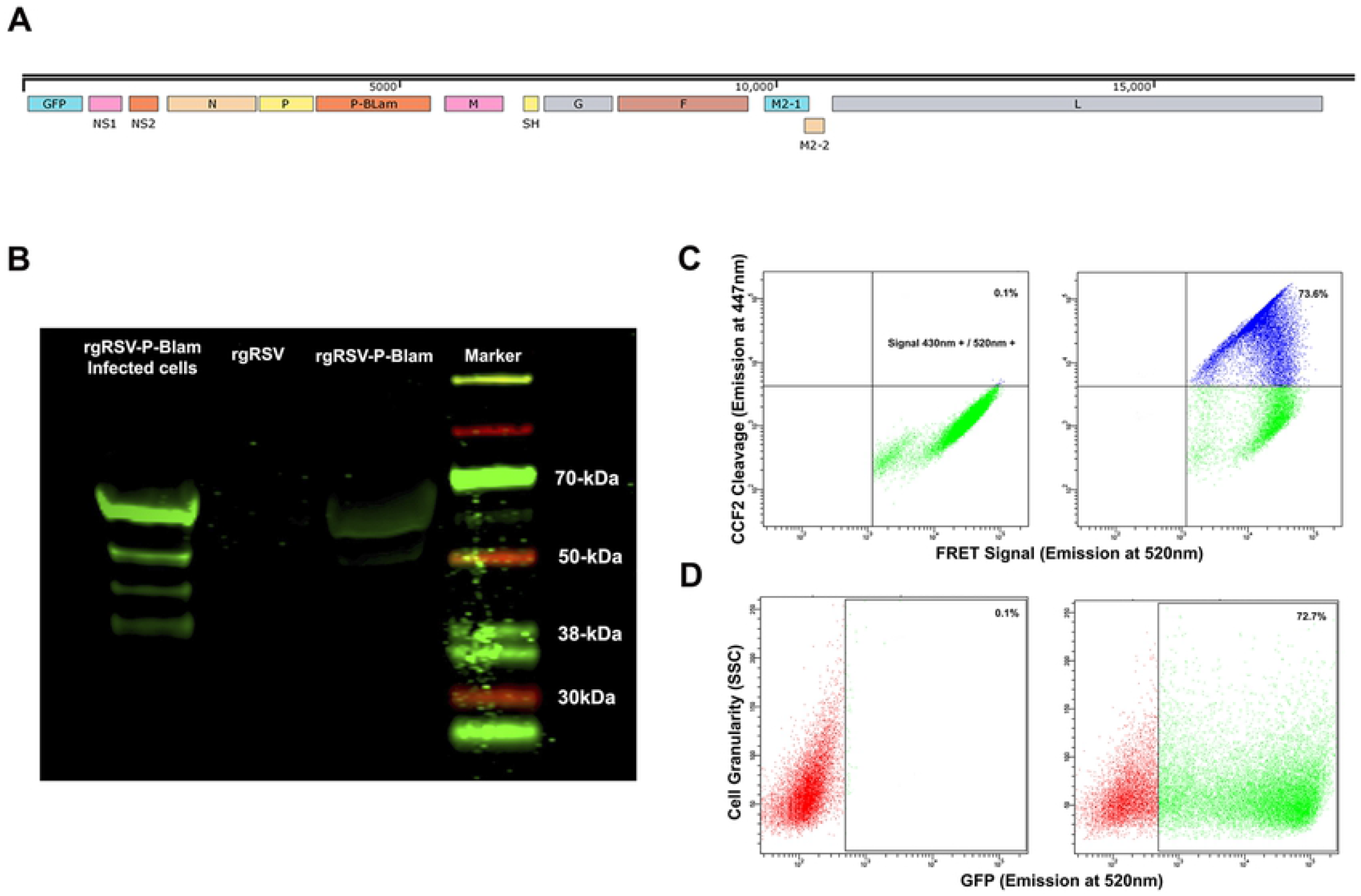
Characterization of rgRSV carrying chimeric protein P-β-lactamase. (A) Position of the P-BlaM gene in the rgRSV-P-BlaM genome. (B) Immunoblot showing recombinant P-BlaM protein in RSV-P-BlaM-infected HEp-2 cell lysates (lane 2); rgRSV virions (lane 3); rgRSV-P-BlaM virions; and MW markers (lane 4). (C, D) Cytometry plots showing the functional activity of the expressed chimeric protein P-BlaM on its CCF2 substrate assessed in NHBE cells infected with rgRSV-P-BlaM (C) along with the assessment of the GFP expression in the same cultures (D). In (C), in the absence of P-BlaM, the FRET mechanism in CCF2 was present revealed by the emission at 520 nm after excitation at 405 nm. In the presence of P-BlaM, CCF2 was cleaved and the FRET was lost allowing the detection of the emission at 447 nm after excitation at 405 nm

For the AB approach, palivizumab was used to neutralize rgRSV-P-BlaM virions attached to the plasma membrane. In NHBE cells, 200 µg/mL of palivizumab was chosen as the working concentration since it blocked 99.7% or 96.6% of infection with rgRSV-P-BlaM in assays probing neutralization of the virions in suspension (**Fig 3A**) or after their attachment to the plasma membrane (**Fig 3B**). Independently of the experimental conditions, such concentration also neutralized infectious doses of MOIs at 0.5 and 2 (**Fig 3**).

**Fig 3.**
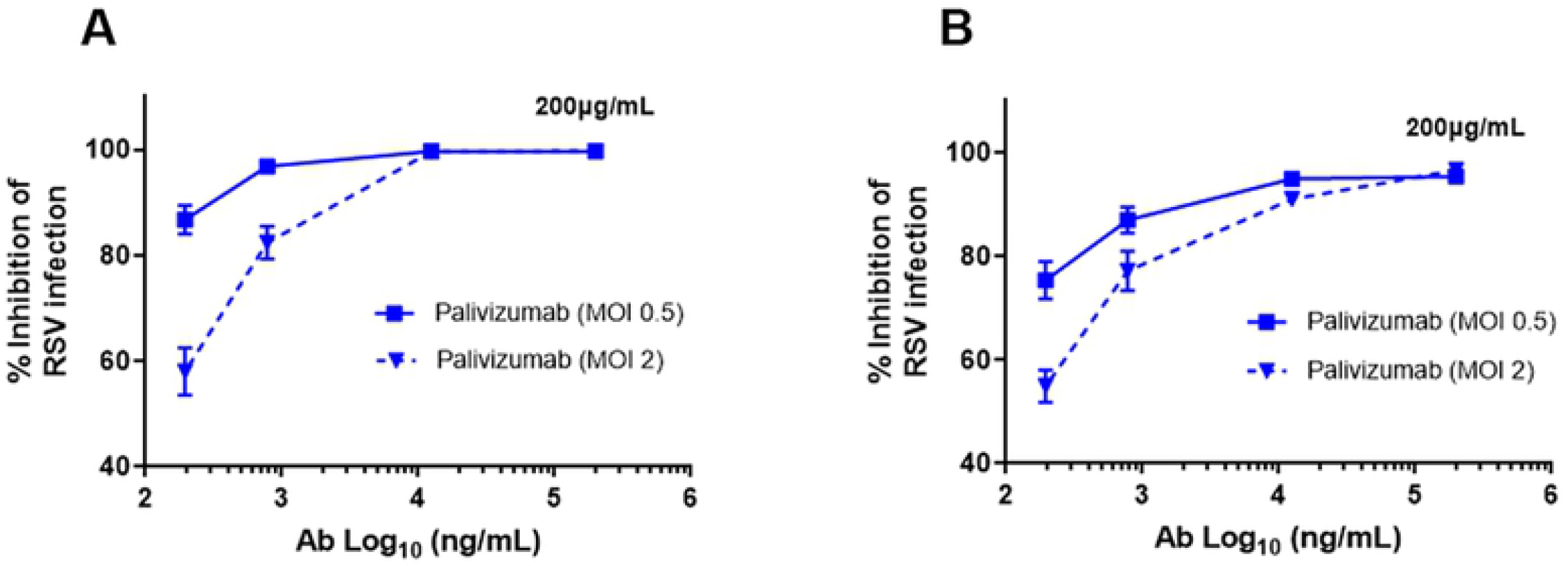
Palivizumab concentration required to neutralize RSV. (A, B) The following concentrations of palivizumab were tested: 200, 12.5, 0.78, 0.195, and 0.048 µg/mL and their Log_10_ transformation were plotted on the X-axis. Two infectious doses of RSV were tested (0.5 and 2.0 MOI). (A) RSV aliquots were incubated with the palivizumab at the aforementioned concentrations for 1 hr at 37°C before adding to the NHBE cultures. (B) RSV aliquots were prebound to NHBE cells for 1 hr at 22°C followed by the addition of palivizumab at the mentioned concentrations. Infection proceeded for 16 h followed and fluorescence-based flow cytometry using GFP expression was used to identify rgRSV-infected cells. Results are mean ± SEM of three independent experiments.

To compare virus entry into NHBE cells following virus attachment at 4°C and 22°C, cell cultures were shifted to 37°C. At the indicated times, virus entry was stopped using either the AB or TB. After 2 hr, the media in all wells was changed to HBSS supplemented with palivizumab (200 µg/ml) and CCF2-AM, the **BlaM** prosubstrate, and incubated for 3 h at 17°C to allow CCF2-AM to enter the cells (**Fig 4A**). Synchronization due to virus binding at 4°C displaced the TB kinetic curve to the right relative to the AB curve (**Fig 4B**), indicating that the attachment at 4°C results in endocytosis-dependent virus entry. However, attaching RSV at 22°C (**Fig 4C**) resulted in similar kinetics for both the AB and the TB assays, indicating that there was no lag between the loss of antibody sensitivity and the nucleocapsid reaching the cytoplasm to begin transcription. That is, fusion with the plasma membrane occurred simultaneously with escape from antibody neutralization. Consequently, the temperature at which virus attachment was conducted biased the route used for RSV to deliver its ribonucleoprotein complex to cytoplasm.

**Fig 4.**
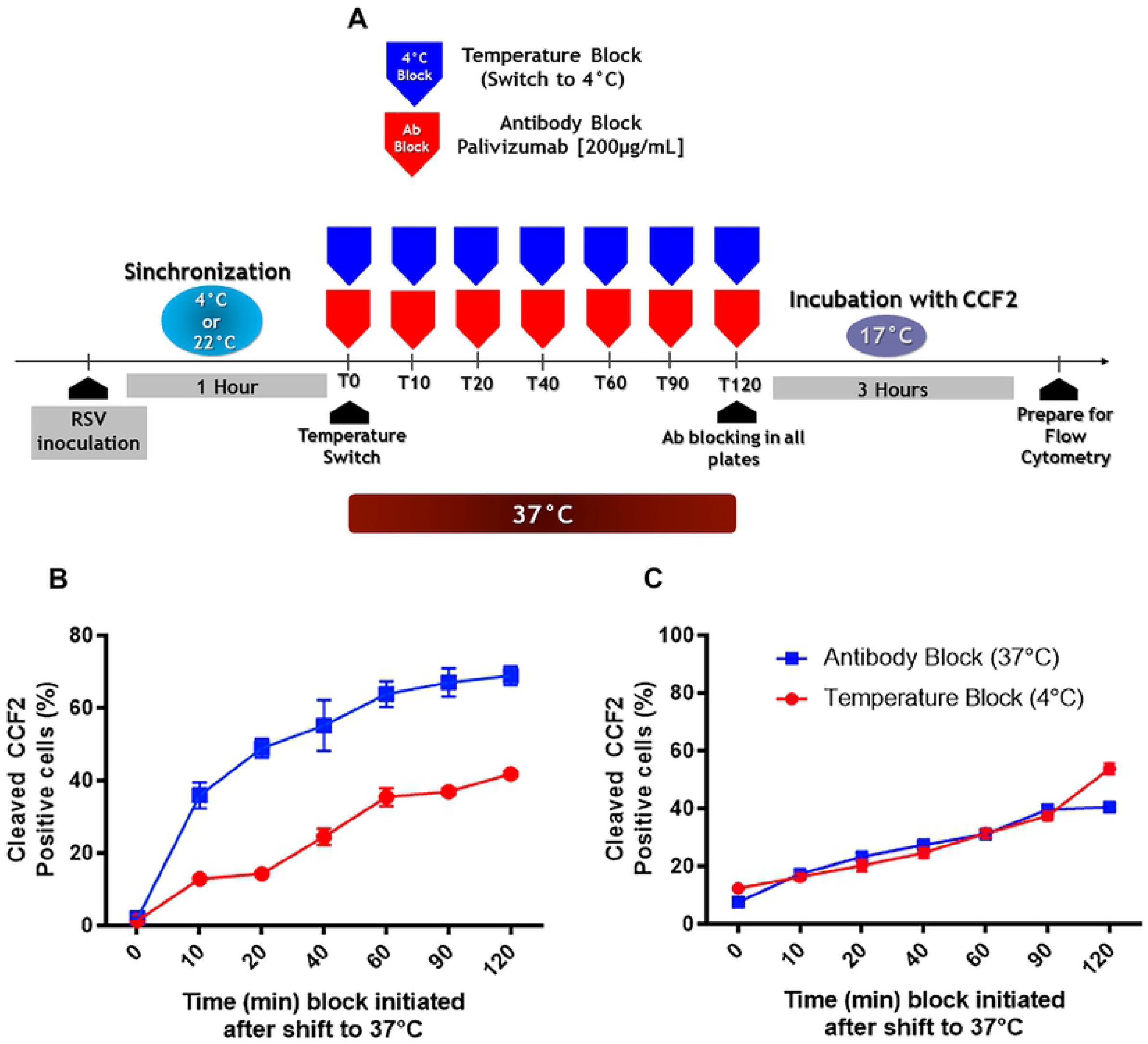
RSV preferred pathway to enter NHBE cells takes different routes depending on the temperature at which attachment took place. (A) Scheme describing the approach that was followed to determine the route that virus used to enter cells according to the temperature at which the attachment was conducted. The attachme5nt of rgRSV-P-BlaM virions proceeded at either 4°C (B) or 22°C (C) for 1 h on NHBE cell cultures, followed by a switch to 37°C. At various times, we stopped virus entry by either antibody block (palivizumab, 200 µg/mL) or temperature block (4°C), washed the cells and loaded them with CCF2-AM at 17°C for 3 h. The plots showed the percentage of cells positive for CCF2 cleavage due to BlaM at each time. Results are mean ± SEM of three independent experiments.

As it is possible that some fraction of the cells was infected by virions entering through endocytosis, we conducted a pulse-chase assay to determine how many of the infected cells were due to virions located in the endosomal compartments. Following the general approach designed by Miyauchi et al [25], a pulse of palivizumab (200 µg/mL) is given in a single dose to all plates 10 min after switching to 37°C from 22°C. Then, the TB-chase consists of placing a set of three plates at 4°C immediately at the moment of antibody administration or at a later time up to 2 hr (**Fig 5A**). Virions that had been endocytosed before the addition of palivizumab would have contributed to the infection, the TB-Chase curve would show an increasing percentage of cleaved-CCF2 positive cells over time, whose curve would be displaced to the right. If virions only entered via the plasma membrane; then, the TB-Chase curve would remain flat in each time point because all virions would be susceptible to palivizumab neutralization. These two possibilities are schematically depicted (**Fig 5B**). The actual experiment (**Fig 5C**) revealed a flat TB-Chase curve compared with the antibody block curve used as a reference, clearly indicating that the vast majority of the virions prebound at 22°C infected by fusing their envelope with the plasma membrane once the temperature was switched to 37°C.

**Fig 5.**
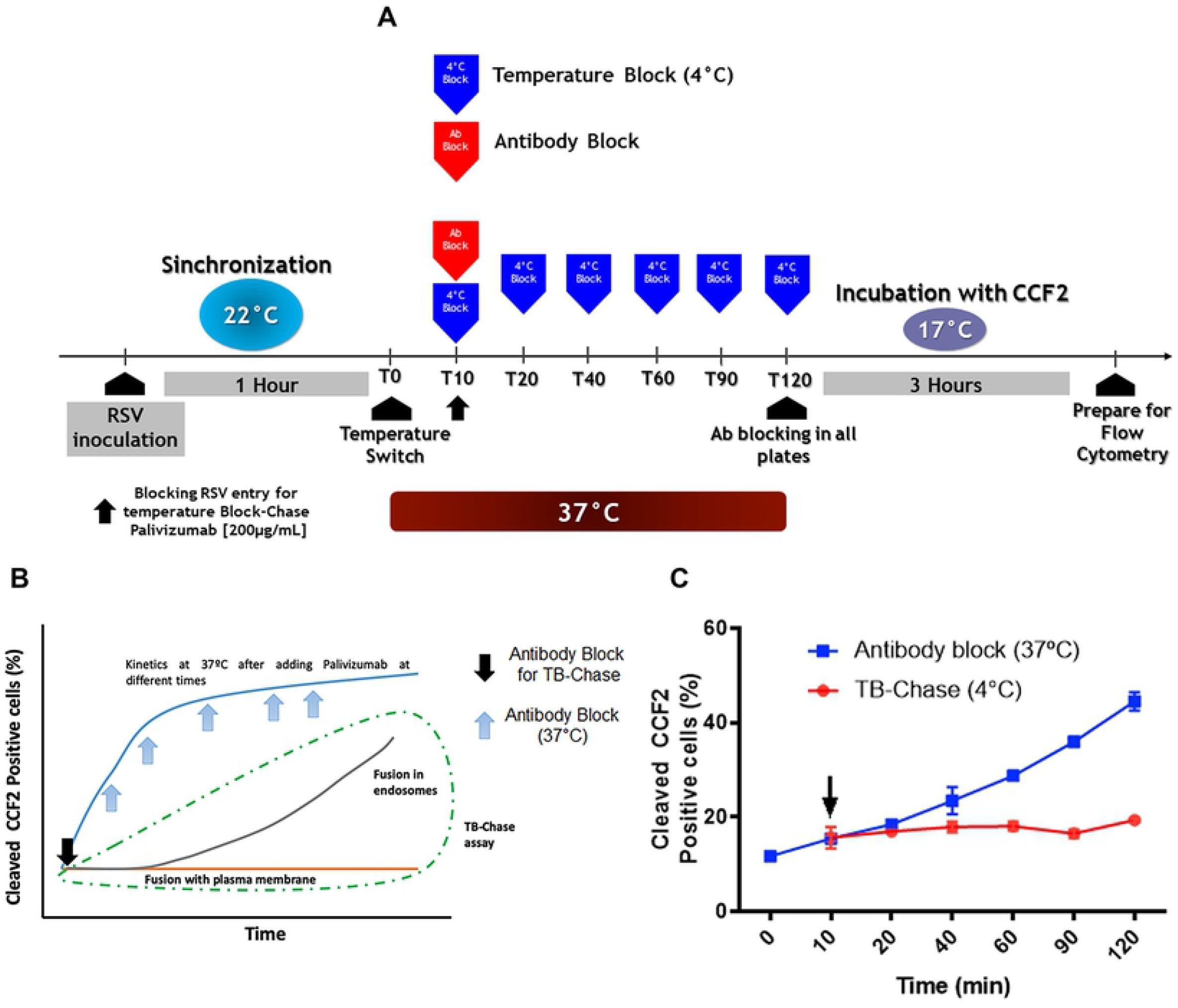
TB-chase assay confirms that the virus entry at 37°C occurs via plasma membrane after attachment at 22°C. (A) Scheme describing the approach that was followed for TB-chase assay that allowed us to determine whether virions inside endosomes contribute to infection. The kinetics at 37°C was assessed at the same times as show in Fig 4. In (B), a schematic drawing shows the different scenarios expected for a TB-chase. The black arrow indicates the time at which we added palivizumab to a set of cell cultures 10 min after switching to 37°C from 22°C. Entry kinetics built from adding palivizumab at different times serves as reference (blue line). In (C), we show the results from the real assay. Here, a black arrow also indicates the time of palivizumab addition to the plates in the TB-Chase. The CCF2-AM loading in the presence of palivizumab proceeded as described in the Fig 4 and Materials and Methods. In the plot, the TB-chase results in a flat line, indicating that RSV virions inside endosomes had almost no contribution to infection after attachment at 22°C. Results are mean ± SEM of three independent experiments.

## Discussion

Overall, the results of this study indicate that the pathway that RSV uses to enter primary cultures of undifferentiated HBEC cells is dependent on the temperature at which attachment was conducted. If the attachment of proceeded at 22°C, RSV enters the target cell by fusing its envelope directly with the plasma membrane. In contrast, synchronization due to virus adsorption at 4°C leads to the erroneous conclusion that virus entry occurs via endosomes. The actomyosin contraction-caused subsidence of the blebbing observed after switching to 37°C from 4°C may be a contributing factor in taking the previously adsorbed virions into the cell.

In this study, we inserted the sequence coding BlaM in RSV genome as a second P gene chimera, P-Blam. Cavrois and colleagues [28] were the first to use beta-lactamase in a virus as part of the BlaM-Vpr chimera to monitor the delivery of HIV-1 content to the cytoplasm. This approach has enabled the discovery of both the mechanism by which HIV-1 infects cell lines and primary cells (T lymphocytes and macrophages) and the role of other cell molecules involved in the viral fusion process [25,29–33]. In addition to HIV-1, BlaM-Vpr was also used to study the entry process of other viruses in models that use pseudotyped lentiviruses carrying envelope proteins derived from either the Hepatitis C virus and Ebola virus [34–36], or in VLPs constructed for influenza A and henipavirus [37,38].

The P-BlaM-associated signal indicates viral content delivery to the cytoplasm. The detection of the P-BlaM protein in the virus preparation as well as the rapid kinetics of the CCF2 cleavage strongly suggested that RSV virions incorporated P-BlaM as a structural protein. Alternatively, or in addition, the nucleocapsid may begin to produce mRNAs from its genome template soon after entry. Either way, our interpretation is the same: the virus has released its content into the cytoplasm.

Incubating the cultures at 4°C for 1 h is the traditional way to synchronize the subsequent fusion step, which facilitates the study of how viruses and cells interact with each other and the cell’s response to infection. However, the fact that incubating at 4°C induces changes in the cytoskeleton and the subsequent formation of actomyosin rings on the plasma membrane when temperature was raised to 37°C may bias the actual course of events that happen during the early steps of virus-cell interaction. On the other hand, there were no obvious cytoskeletal changes at 22°C, even though this temperature is considered suboptimal because it slows down cellular processes and membrane fusion.

Adsorption at 4°C causes the virus to enter by endocytosis once the temperature is switched to 37°C as demonstrated by the shift to the right in terms of the cumulative BlaM positive events seen in the condition of temperature block compared with antibody block. Such endocytosis was previously associated to macropinocytosis [39] and may be a consequence of the blebbing as we have shown that incubation at 4°C causes blebbing. On the other hand, virions attached at 22°C delivered their ribonucleoprotein complex to the cytoplasm via fusion at the plasma membrane once the inhibition of membrane fusion was reversed at 37°C. Consequently, entry via macropinocytosis is an artefact. It is not a normal cell response triggered by the virus.

The fact that attaching virions at 4°C induced endocytosis as a cellular response partially explains previous reports that have reported RSV entry via macropinocytosis [20,23]. Consequently, it may also be necessary to reconsider reports of other viruses whose main conclusion was that endocytosis is their preferred entry pathway since the experiments that led to that conclusion used attachment at 4°C to synchronize virus infection [40–50].

Although drugs whose mechanism of action have been used to suggest macropinocytosis [20,23,51], their pharmacological effect may also have off-target effects that may obscure the real mechanism or conduct to erroneous interpretation. Using siRNAs library targeting cell transcripts have provided contradictory results suggesting RSV entry dependent on either clathrin- or macropinocytosis-mediated pathways [22,52]. Our approach using a temperature closer to 37°C for the attachment step as well as a system that rapidly detects virion content delivery was key to determining the entry route of RSV via the plasma membrane in primary, though not differentiated, cultures of human bronchial epithelial cells. By tracing MTRIP-labeled RSV in real-time, Alonas et al [19] found that RSV passively moves on the cell membrane, followed by transient rapid movement and changes in the virion’s shape. They proposed that such behavior was akin to RSV fusing its envelope with the plasma membrane because of the assumption that endosomes must show a dynein-dependent active transport [19].

Histologically, the bronchi are lined by pseudostratified columnar ciliated epithelium [53], meaning that although all cells have contact with the basal membrane, not all of them are exposed to the lumen. We used normal human bronchial epithelial cells grown in a monolayer, which in the presence of ROCK inhibitor possesses characteristics of basal cells [24]. These basal cells differentiate into cells that have access to the lumen, such as ciliated, goblet, and pulmonary ionocytes [53–55]. Immunohistopathological evaluation of lung tissues derived from autopsies of children who died from RSV infection showed that ciliated cells are the most likely ones to be infected by this virus [56,57]. This tropism has also been observed in well-differentiated airway epithelial cultures [58,59]. However, mechanical or chemical damage of well-differentiated respiratory epithelial cultures exposes basal cells, which could consequently be infected by RSV and promote the virus to spread throughout the epithelium [60]. Thus, our results may apply to patients whose airways are affected by diseases such as asthma or COPD – patients who have a hyperproliferative basal epithelium and a malfunctioning epithelial barrier due to an alteration of tight junctions [61]. Future studies should be aimed at evaluating viral entry in well-differentiated respiratory epithelial cultures.

In summary, we have found that RSV infects primary basal bronchial epithelial cells by fusing its envelope with their plasma membrane. However, synchronizing infection by cooling the cells to 4°C for the inoculation period changed the entry mechanism, resulting in endosomal entry. This problem can be avoided by using a higher adsorption temperature for synchronization, such as 22°C, that does not significantly affect cellular dynamics while at the same time avoids triggering the fusion mechanism.

## Materials and Methods Cell culture

We acquired undifferentiated normal human bronchial epithelial (NHBE) cells from Lonza, while murine fibroblasts – 3T3-J2 – were purchased from Kerafast. We used conditionally reprograming for a continuous expansion of NHBE cells in order to take the number of passages beyond the critical point of senescence [62]. For such conditional reprogramming, undifferentiated NHBE cells were grown on a bed of Mitomycin-treated 3T3-J2 fibroblast cells in the presence of the ROCK inhibitor, Y27632 (Tocris).

For 3T3-J2 cell expansion, we cultured them in 3T3 medium: DMEM (ThermoFisher Scientific) supplemented with 10% bovine calf serum (ThermoFisher Scientific) and 1% penicillin-streptomycin (ThermoFisher Scientific). Cells were incubated at 37°C with 5% CO_2_, and 95% relative humidity until confluency reached 70%. Cells were then seeded in new T-175 flasks (Corning) at a density of 3.5×10^3^ cells per cm^2^. After 80h, cell monolayer was washed with 1X PBS followed by adding fresh 3T3-J2 media supplemented with 2 μg/mL Mitomycin C (Sigma Aldrich) to inactivate mitosis. After removing Mytomicin-supplemented 3T3 medium, we carefully washed the cell monolayer with 1x PBS thrice to remove Mitomycin excess. Then, cell monolayer was detached from the surface using Trypsin (Sigma Aldrich). After trypsin neutralization and cell washing and pelleting by centrifugation, cells were seeded at a density of 2×10^4^ cells/cm^2^ in T75 or T175 flasks coated with collagen. After overnight incubation for cell adhesion, the Mytomicin-treated 3T3-J2 cells were ready to support the expansion of epithelial cells.

Primary undifferentiated NHBE cells were seeded on Mytomicin-treated 3T3-J2 bed in F medium supplemented with the ROCK inhibitor Y-27632 (7.5 µM), as previously described by Liu *et al*. [63]. Cultures were kept at 37°C with 5% CO_2_ and 95% relative humidity. NHBE cells became to displace 3T3-J2 and at the same time formed round colonies. Once they reached 70% confluence, we incubated the culture in PBS + EDTA (Dulbecco’s 1×PBS; 100 mM EDTA) for 5 min to detach 3T3-J2 fibroblasts. Then, NHBE cells were detached using a Trypsin-based commercial kit from Lonza following respective recommendations. Finally, NHBE cells were cryo-preserved in F medium supplemented with 10% FBS, 10% DMSO and 7.5 μM ROCK inhibitor.

For experimental assays, the undifferentiated NHBE cells were cultured and expanded in BEGM medium (Lonza) supplemented with 7.5 μM ROCK inhibitor in T75 flasks for 3 days. Then, NHBE cells were sub-cultured and seeded in 24-well plates at a density of 1.2×10^5^ cells/well in the presence of ROCK inhibitor. After overnight incubation, we removed Y27632 by changing medium. After 24h incubation in BEGM, cells reached 80% confluency and we used them for viral infection assays.

### Assaying F-actin state by staining with Alexa-488-labeled phalloidin

We evaluated whether temperature may have an impact on the actin cytoskeleton by staining F-actin with phalloidin. Briefly, NHBE cells were subcultured on rounded polylysine-coated glass coverslips (PDL, Neuvitro). At 70% confluency, cell cultures were incubated at different temperatures (4°C, 22°C or 37°) for 1 h. A different set of cultures were used to explore the effect of a sudden temperature change on F-actin cytoskeleton. This was conducted by exposing cultures to either 4°C or 22°C, which was then followed by subsequent incubation at 37°C for 10 min. After temperature treatment, we fixed NHBE cell cultures with 4% paraformaldehyde for 15 min at 4°C, which was followed by permeabilization with Triton X-100 for 15 min a room temperature. Finally, we incubated each cell culture with Alexa-488-labelled phalloidin (ThermoFisher Scientific) in order to tag F-actin. We used FluorSafe mounting medium (Calbiochem). The cells were visualized at 63X (NA 1.4) using a confocal microscopy Axio observer Z1. Using Zen-Blue software, multiple Z-plane frames were serially acquired with a separation of 0.5 µm from each other along the Z axis using the module Z-stack. Then, the images were merged and subjected to signal deconvolution using the module Extended Depth of Focus.

### Design, construction and production of rgRSV-P-BlaM

We have the sequence encoding the chimera P-BlaM made by integrated DNA Technologies as gBlocks. P stands for the RSV phosphoprotein, and BlaM corresponded to the optimized beta lactamase version (Y105W) [38]. By Gibson cloning (New England Biolabs), we inserted the gBlock at the sequence flanked by AvrII (nt. 2930) and PvuI (nt. 6573), using HC123 full-length RSV cDNA construct as a backbone. The HC123 was derived from cDNA 224 [64]. The modified HC123 cDNA construct sequence has P-BlaM positioned after the RSV-P gene.

We used BHK cells constitutively expressing T7-RNA polymerase (BHK/T7 cells) to make rgRSV-P-BlaM virions. BHK/T7 cells were subcultured in 6-well plates fed with DMEM-Glutamax medium (ThermoFisher Scientific) supplemented with 2% FBS (ThermoFisher Scientific) and 1% Penicillin-Streptomycin (ThermoFisher Scientific). At 70% confluency, BHK/T7 cells were transfected with P-BlaM RSV cDNA and helper plasmids [65] using Lipofectamine 3000 (ThermoFisher Scientific) – in the following quantities: 800 ng of modified HC123 cDNA vector, 400 ng of N, P, and M2-1, and 200 ng of L. The transcription in each of the constructs were under the control of the T7 promoter. Following 2 days, transfected cells were subcultured in T25 flask (Corning), and syncytia formation was monitored for 48 h. This was then followed by a subsequent subculture in T75 flask (Corning). We proceeded to change medium each 48 h until we observed around 50%-60% syncytia in the monolayer. Cells were then scraped in a small fraction of the supernatant. Both scraped cells and supernatant were placed in the same Falcon tube, followed by a subsequent centrifugation at 2,141 g for 10 min at 4°C to pellet the cell debris. Magnesium sulfate at 0.1 M was used to stabilize the virus suspension. We snap-froze 1-mL aliquots in dry ice. Each aliquot was stored at −150°C in a cryofreezer (Panasonic).

### Production and titration of recombinant virions (rgRSV-P-BlaM and rgRSV)

We grew recombinant respiratory syncytial virus (RSV) in HEp-2 cell cultures. Briefly, HEp-2 cells were expanded in Opti-MEM™ medium supplemented with Glutamax (ThermoFisher Scientific),10% FBS (ThermoFisher Scientific) and 1% Penicillin-Streptomycin (ThermoFisher Scientific) in T75 flasks for 3 days at standard culture conditions of 37°C, 5% CO_2_, and 95% relative humidity. Subsequently, cells were transferred to T175 flasks at such cell density to reach 60% confluency next day. After washing with Ca^++^- and Mg^++^-free PBS, HEp-2 was infected with rgRSV at a multiplicity of infection (MOI) of 1 infectious viral particle per 10 cells (MOI of 0.1). The infection proceeded in Opti-MEM (ThermoFisher Scientific) supplemented with 2% FBS during 48 h while monitoring for cytopathic effects and the formation of syncytia. At such time, we changed medium and left the culture proceed for 20 h more. Then, we transferred approximately ¾ of the medium to 50-mL Falcon tube. We scraped the cells in the medium that was left in each flask and transferred them to the abovementioned 50-mL Falcon tube. The cell debris was pelleted by centrifugation at 2,141 g at 4°C for 10 min in a Sorvall Legend Match 1.6 centrifuge. The infectious supernatant was transferred to a new pre-chilled, labelled Falcon tube. We supplemented the supernatant to stabilize the virions with the following compounds at the indicated final concentration: 0.1 M MgSO_4_ (Sigma Aldrich), 0.1% human Albumin (Biotest) and 50 mM HEPES (Gibco). One-mL aliquots of supernatant in cryotubes were snap-frozen in dry ice. Subsequently, these were stored in a −150°C freezer (Panasonic).

We estimated the virus titer by preparing five-fold serial dilutions from stock in HEp-2 cell cultures, which were previously subcultured in 24-well plates. After standard infection for 2 h at 37°C, infection was allowed to proceed for 16 h at 37°C. Trypsin treatment was used to detach cells from the well surface. Using fluorescence-based flow cytometry, we detected RSV-infected cells. Following Techaarpornkul et al [66] recommendations, we calculated the viral titer from that dilution that gave a MOI in the range of 0.05 to 0.1. The formula to calculate such MOI is: (% of infected cells × total number of cells × dilution factor) / (100 × volume of the infection aliquot)

### Western blot assay to demonstrate the presence of P-BlaM

The processing of rgRSV-P-BlaM-infected HEp-2 cells to obtain the respective lysates consisted of scraping in cold lysis buffer supplemented with protease inhibitor (10 µL/mL). After centrifugation, the supernatant is stored at −80°C until further use. We pelleted the rgRSV-P-BlaM virions by centrifuging the viral preparations at 20,000 rpm for 2 hr at 4°C. After resuspending the pellet in 100 µL of lysis buffer supplemented with protease inhibitor, it was boiled for 5 min. Then, samples were processed and run in SDS-PAGE under reducing conditions. After blotting, the primary antibody used to probe for the presence of P-BlaM was mouse monoclonal anti-BlaM antibody [clone 8A5.A10] (ABCAM, ab12251). The 800cw goat anti mouse IgG1 from Odyssey near-infrared system (LI-COR, Lincoln, Nebr., USA) was the secondary antibody used to visualize the primary antibody binding to BlaM.

### Assessment of palivizumab concentration to neutralize RSV

We tested a range of concentrations to determine the neutralization profile of palivizumab (AbbVie). Virus and antibodies were incubated together in solution for 1 h at 37°C before adding the mixing to NHBE cell monolayer. We tested the following antibody concentrations in µg / mL: 200, 12.5, 0.78, 0.195, and 0.048. These antibody solutions were prepared in BEGM medium. RSV infection was allowed to proceed for 16 h at 37°C. Then, cell cultures were processed for fluorescence-based flow cytometry. Infected cells were identified by GFP expression.

### Assays for determining RSV entry kinetics after synchronizing viral entry by adsorption at different temperatures (4°C or 22°C)

After washing with HEPES-based saline solution (HBSS), NHBE cells at 80% confluency were infected with rgRSV-P-BlaM at different infectious doses (please see respective results section) suspended in BEBM. The virus adsorption proceeded at either 22°C or 4°C for 1 h to synchronize viral fusion.

After synchronization at the chosen temperature, plates were placed in the incubator at 37°C. At different time intervals (0, 10, 30, 45, 60, 90 and 120 min), we stopped the infection either by using palivizumab (antibody block) or by placing a selected plate at 4°C (temperature block - TB).

When we used BlaM activity on CCF2-AM as a way to identify RSV-infected cells, we stopped the infection by either antibody (200 µg / mL palivizumab) neutralization or TB (4°C) treatment at the respective interval. When the last time interval was reached (150 min), we changed the rgRSV-P-BlaM-containing medium for HBSS supplemented with 2 µM CCF2-AM and 200 µg / mL palivizumab in all culture plates. CCF2-AM (GeneBLAzer in vivo Detection Kit, ThermoFisher Scientific) was allowed to be loaded into cells by incubating each culture for 3 h at 17°C. Finally, cells were prepared for fluorescence-based flow cytometry. CCF2 is a compound made up of a hydroxycoumarin moiety linked by a beta-lactam ring bridge to fluorescein. In the absence of BlaM, FRET mechanism is present and is determined by measuring emission at 520 nm after excitation at 405 nm. In the presence of BlaM, the FRET mechanism is lost as both molecules are separated by catalytic cleavage; consequently, we detected the emission at 447 nm after excitation at 405 nm.

We also ran a TB-chase assay in which after synchronizing rgRSV-P-BlaM adsorption at 4°C for 1 h, we added 200 µg / mL palivizumab to each culture when switching the temperature to 37°C. After each abovementioned time interval, we placed the respective plate at 4°C to prevent virus entry at endosome. When the last time interval was reached (150 min), cells were processed as abovementioned for determining BlaM activity using CCF2-AM.

When we used GFP fluorescence as readout, we stopped the infection by changing the rgRSV-containing medium for BEGM supplemented with the respective antibody (either 200 µg / mL palivizumab or 1.5 µg / mL D25) to neutralize active virions still adsorbed to the plasma membrane. Then, we allowed the infection to proceed for 16 h before preparing the cells for fluorescence-based flow cytometry (BD FACS CANTO II).

## Acknowledgement

Research reported in this publication was supported by the National Institute of Allergy and Infectious Diseases of the National Institutes of Health under Award Number R01AI110385 (HS) and R01 AI093848 (MEP). The content is solely the responsibility of the authors and does not necessarily represent the official views of the National Institutes of Health. HS was also partially supported by a Fulbright-Universidad del Norte Research visiting scholarship.

